# Chromatic bacteria v.2 - A Himar1 transposon based delivery vector to extend the host range of a toolbox to fluorescently tag bacteria

**DOI:** 10.1101/2021.03.26.437257

**Authors:** Christian Stocks, Rudolf Schlechter, Mitja Remus-Emsermann

## Abstract

A recent publication described the construction and utility of a comprehensive “Chromatic Bacteria” toolbox containing a set of genetic tools that allows for fluorescently tagging a variety of Proteobacteria. In an effort to expand the range of bacteria taggable with the Chromatic Bacteria toolbox, a Himar1 transposon vector was constructed to mediate insertion of fluorescent protein and abiotic resistant genes. The Himar1 transposon was chosen as it is known to function in a wide range of bacteria. To test the use of the Himar1 derivative of the Chromatic Bacteria plasmid series for fluorescently tagging bacteria, recently isolated non-model organisms were conjugated with the Himar1 transposon plasmids. We were unsuccessful in delivering the plasmids into Gram positive bacterial isolates but were able to successfully integrate the transposon in isolates that we were previously unable to modify such as *Sphingomonas* sp. Leaf357 and *Acidovorax* sp. Leaf84. This manuscript reports on the currently available plasmids and transposition success in different bacteria. The manuscript will be updated regularly until a full plasmid series is available and a large range of recipient strains was tested.

## Introduction

Recently we described the constructions of a comprehensive “Chromatic Bacteria” toolbox containing a set of plasmids denoted the pMRE series (Schlechter et al., 2018). These tools can be used to fluorescently tag a wide range of bacterial isolates, however, there were several isolates that were not able to genetically modify. Fluorescently tagging bacteria using genetic manipulation is a state of the art technology to track bacteria *in situ* in *in vitro* to study their behaviour such as on plant leaf surfaces (Nadell et al., 2016; Remus-Emsermann and Schlechter, 2018; Bernach et al., 2019; Miebach et al., 2020).

By fluorescently tagging bacteria, it is possible to study them at the micrometer resolution. This is in stark contrast with the meta-omic research which has driven microbiology and microbial ecology during the last decade. Fluorescent tagging allows the study of bacteria at single cell resolution, which gives insight into biofilm formation, bacteria-bacteria and bacteria-host interactions (Schlechter et al., 2018). The recently published Chromatic Bacteria toolbox employs three different vectors for fluorescent tagging, paired with one of eight fluorescent proteins and additionally one of four combinations of antibiotic resistant cassettes. The three different vectors are based on i) a broad-host plasmid ii) a Tn*5* transposon delivery plasmid and iii) a Tn*7* transposon delivery plasmid. We have determined the host-range of the Chromatic Bacteria toolbox by extensive plasmid conjugation experiments. As a result, it was shown that, even though wide, the utility of the toolset’s host range is limited to Proteobacteria (Schlechter et al., 2018).

To further expand the host range of the Chromatic toolbox, we expanded the toolset by incorporating a Himar1 transposon-based delivery system. The Himar1 transposable element belongs to the mariner family of transposons. Mariner transposons can be found throughout eukaryotic and prokaryotic organisms (Lampe, 2010; Tellier et al., 2015). The Himar1 transposon gene was discovered in a horn fly and has been mutated to construct a hyperactive version (Lampe et al., 1996). The gene encodes for a transposase protein that functions through a cut and paste mechanism and cleaves sequences between Thymine Adenine (AT) dinucleotide sites. Unlike other transposases such as the Tn*5* transposase, the Himar1 transposase requires no cofactors to initialise or enable gene transposition that need to be provided *in trans* by the host organism (Lampe et al., 1996). These low requirements for transposition make Himar1 ideal for random insertion mutagenesis and gene delivery into organisms that do not provide such cofactors. The Himar1 transposase has been shown to function in Gram negative bacteria such as *Pseudomonas fluorescens* and *Flavobacterium johnsoniae* (Braun et al., 2005; McCully et al., 2019) and Gram positive bacteria including *Staphylococcus aureus, Clostridium perfringens, Streptococcus mutans, and Mycobacterium smegmatis* (Rubin et al., 1999; Li et al., 2009; Liu et al., 2013; Nilsson et al., 2014). In this study we describe the construction and utility of Himar1-based suicide vectors that allow the fluorescent tagging of bacteria.

## Materials and Methods

### Strains and media

All strains, media, and growth conditions used in this study are listed in Table 1. Lysogeny broth (LB, HiMedia), nutrient broth (NB; HiMedia), Reasoner’s 2A media (R2A, HiMedia), and Reasoner’s 2A agar (R2A agar, HiMedia) were produced according to manufacturer’s instructions. Media were supplemented with 1.5% bacteriological agar (Oxoid) where needed. To support growth of *E. coli* ST18, media was supplemented with 50 mg L^−1^ 5-aminolevulinic acid (5-ala, Sigma). When appropriate, media were supplemented with antibiotics with the following concentrations: 100 mg L^−1^ ampicillin, 20 mg L^−1^ chloramphenicol, 10 mg L^−1^ colistin, 100 mg L^−1^ erythromycin, 15 mg L^−1^ gentamicin, and/ or 50 mg L^−1^ kanamycin.

**Table 1:**
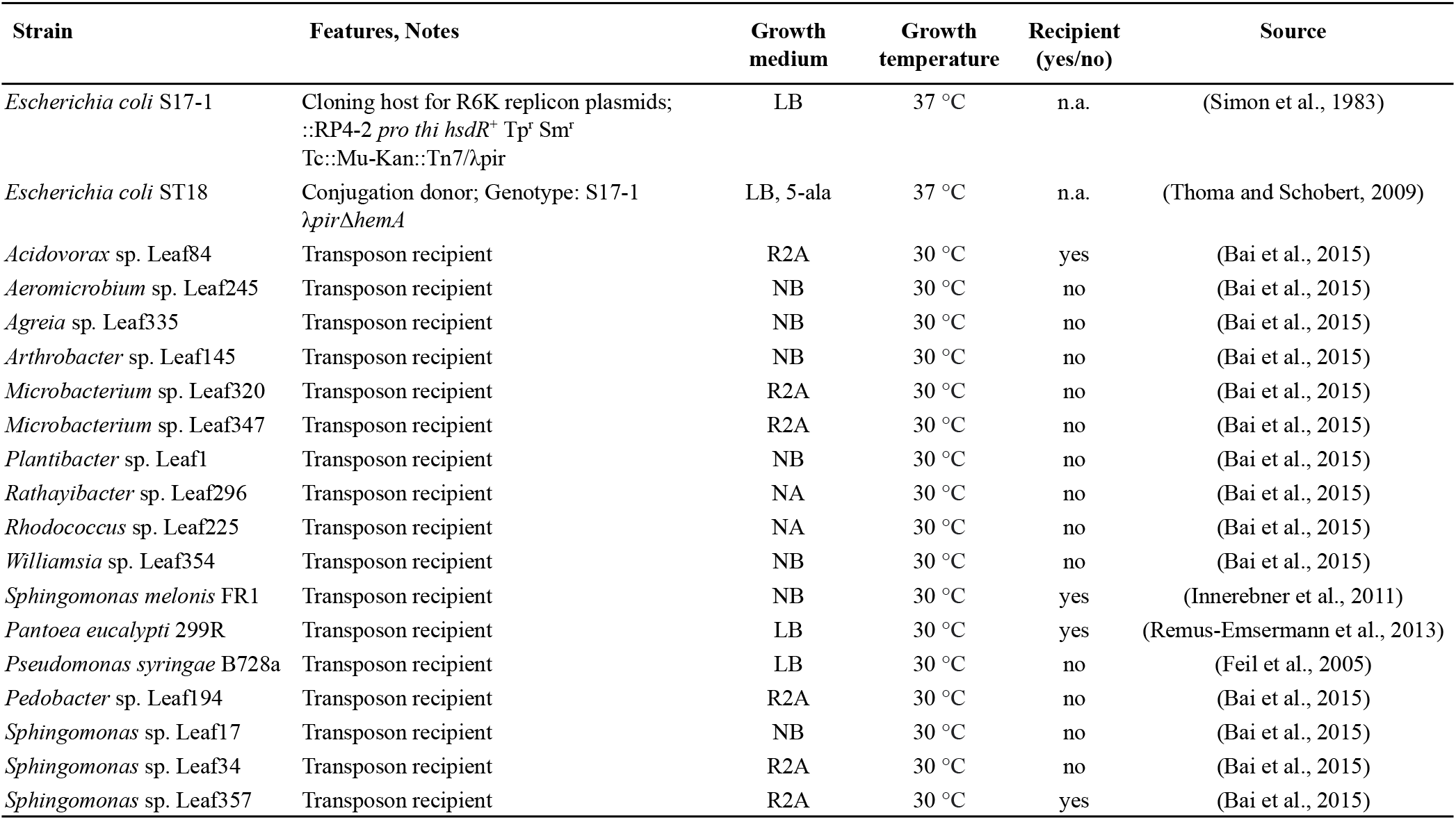
Bacterial strains used in this work.

### Plasmid construction

An overview of the primers and plasmids used in this project are given in Table 2 and Table 3, respectively. Plasmids were constructed as previously described (Schlechter et al., 2018). The here described plasmids are denoted pMRE-Himar-1XY series, where X can either be 3, 4, 5, or 9, representing the different antibiotic resistance gene combinations of chloramphenicol; gentamicin and chloramphenicol; kanamycin and chloramphenicol; or erythromycin, kanamycin and chloramphenicol, respectively, and where Y can be 1, 2, 3, 4, 5, 6, or 7 representing cyan, green, yellow, orange, red, near-infrared or green FP, respectively (mTurquoise2, sGFP2, mOrange2, sYFP2, mScarlet-I, mCardinal, or mClover3, respectively).

**Table 2:**
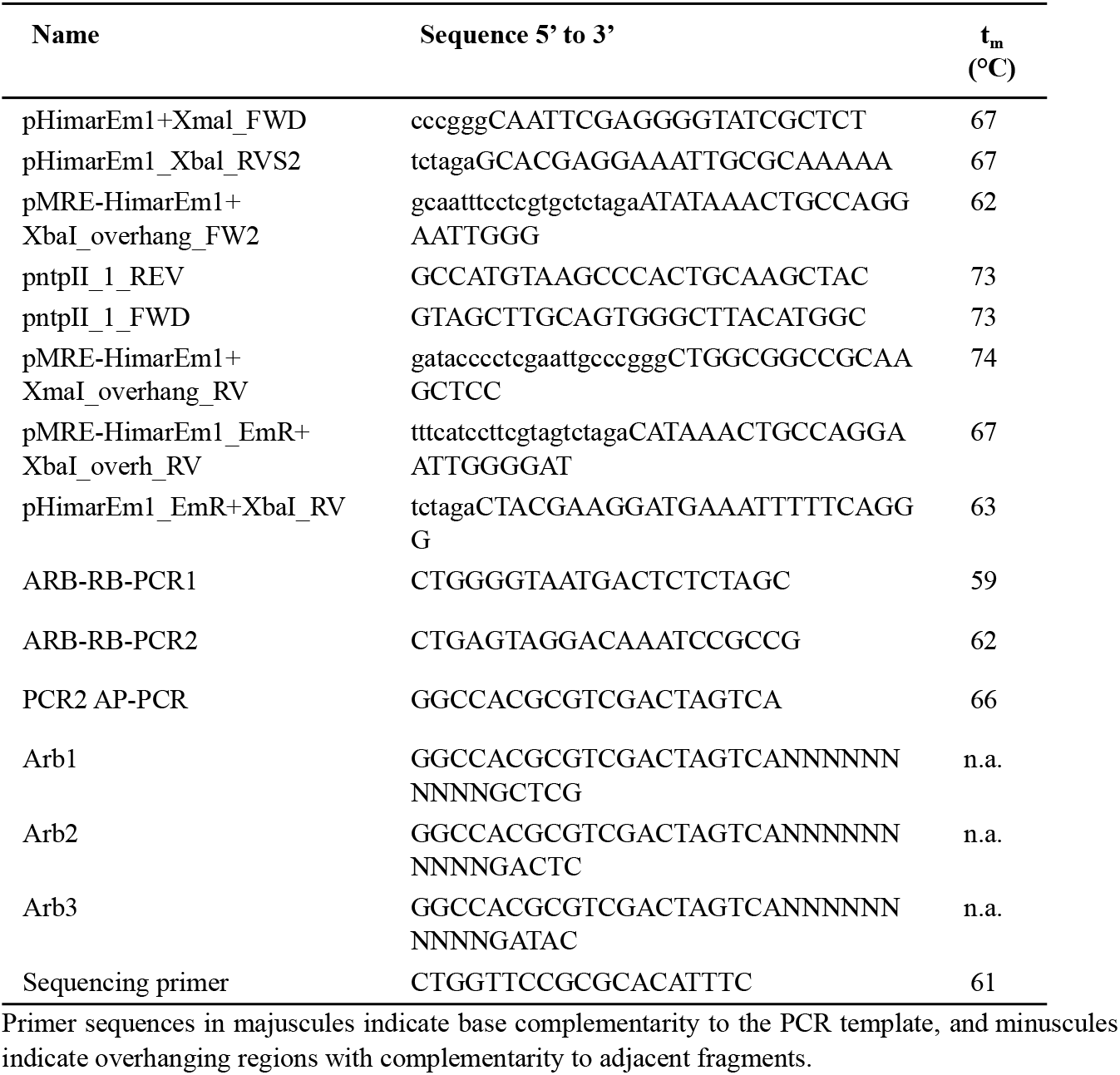
Primers used in this study

**Table 3:** Plasmids used in this study

| Name | Features | Source |
| --- | --- | --- |
| pMRE-13X | Fluorescent protein genes, Cm<sup>R</sup> | Schlechter et al., 2018 |
| pMRE-14X | Cm<sup>R</sup>, Gm<sup>R</sup> | Schlechter et al., 2018 |
| pMRE-15X | Cm<sup>R</sup>, Kan<sup>R</sup> | Schlechter et al., 2018 |
| pHimarEm1 | Himar1 transposon, Kan<sup>R</sup>, Erm<sup>R</sup> | Braun et al., 2005 |
| pMRE-Himar-13X<br>X= 1, 3, 4, 5, 7 | Himar1 transposon, Cm<sup>R</sup> | This work |
| pMRE-Himar-14X<br>X= 1, 2, 3, 4, 5, 6, 7 | Himar1 transposon, Cm<sup>R</sup>, Gm<sup>R</sup> | This work |
| pMRE-Himar-15X<br>X= 1, 3, 5, 7 | Himar1 transposon, Cm<sup>R</sup>, Km<sup>R</sup> | This work |
| pMRE-Himar-19X<br>X= 1, 2, 3, 4, 5, 6, 7 | Himar1 transposon, Cm<sup>R</sup>, Erm<sup>R</sup>, Km<sup>R</sup> | This work |

"X" denotes fluorescent protein genes 0=mTagBFP2, 1=mTurquoise2, 2=sGFP2, 3=sYFP2, 4=mOrange2, 5=mScarlet-1, 6=mCardinal, 7=mClover3.

Each plasmid was constructed by amplifying three fragments: the plasmid backbone, a unique antibiotic resistance gene (X fragment), and a unique FP gene upstream of a chloramphenicol resistance gene (Y fragment). All PCRs were performed using Phusion High-Fidelity Polymerase (Thermo Scientific). For PCR mixes containing primers with overhangs, a touchdown PCR protocol was performed starting with an annealing temperature 10 °C above the *t*_m_. This was reduced by 1 °C for ten cycles before running the PCR with the annealing temperature set to *t*_m_. After amplification, all PCR reactions were DpnI treated to digest the methylated plasmid template DNA before the PCR fragments were purified with the DNA Clean & Concentrator Kit (Zymo). To construct pMRE-Himar-13X, 14X and 15X, the plasmid backbone was amplified from pHimarEm1 using pHimarEm1+XmaI_FWD and pHimarEm1_Xbal_RVS2. Antibiotic resistance genes and fluorescent protein genes were amplified from pMRE-13X, pMRE-14X, or pMRE15X. The X fragments were amplified using primers pMRE-HimarEm1+XbaI_overhang_FW2 and pntpII_1_REV. The Y fragments were amplified using primers pntpII_1_FWD and pMRE-HimarEm1+XmaI_overhang_RV. For construction of the pMRE-Himar-19X series, pHimarEm1 was amplified using primers pHimarEm1+XmaI_FWD and pHimarEm1_EmR+XbaI_RV, which included the pHimarEm1-borne erythromycin resistance gene. An additional kanamycin resistance gene was amplified from pMRE-15X using primers pMRE-HimarEm1+Xbal_overhang_FW2 and pntpII_1_REV. The FP fragment was amplified using the primers pntpII_1_FWD and pMRE-HimarEm1_EmR+XbaI_overh_RV from pMRE-15X plasmids.

Gibson assembly was performed as previously described (Schlechter et al., 2018). Briefly, the fragments were mixed at a 1:3, backbone:insert molar ratio with between 20-100 ng of backbone fragment being used. No more than 5 μL DNA solution was added to 15 μL Gibson assembly mix. Where appropriate, water was added to top up the reaction volume to 20 μL. The Gibson assembly mix was incubated at 50 °C for 20 minutes. Chemically competent *E. coli* S17-1 cells were then transformed using 10 μL of the Gibson assembly mix (Sambrook et al., 1989). After the plasmids were confirmed by Sanger sequencing, they were cloned into chemically competent *E. coli* ST18 (Sambrook et al., 1989).

### Transposon delivery using conjugation

Two parental matings were performed to deliver the pHimar1Em-based plasmids into a range of bacterial strains, following the protocol described by Schlechter and Remus-Emsermann (2019). In variation to this protocol, conjugations were performed using the auxotrophic *E. coli* ST18 as a plasmid donor strain (Thoma and Schobert, 2009). To perform the conjugations, recipient strains (Table 1) were grown in 50 mL of suitable media for up to three days depending on their growth rate. Single colonies of *E. coli* ST18 donor strains were produced and used to inoculate overnight cultures of LB supplemented with appropriate antibiotics and 5-aminolevulinic acid. From these overnight cultures 2 mL were inoculated into 100 mL fresh LB. The *E. coli* donor cultures were grown to an OD_600nm_ of 0.5, and then both donor and recipient cells were collected by centrifugation and resuspended in phosphate buffered saline (PBS) to an OD_600nm_ of 1. Approximately 5 ml recipient and donor cells were next combined in a 1:1 ratio, then centrifuged and resuspended in 200 μL of PBS. The suspension was then spotted onto a 0.44 μm S-pak Membrane filter (Millipore) which was placed on LB agar plates supplemented with 5-ala. The bacterial mixes were incubated at 30 °C overnight. Bacteria were recovered from the filter by vigorous vortexing in 10 mL PBS in 50 mL falcon tubes. Subsequently, the filter was dismissed and the mixes were concentrated by centrifuged and resuspension in 1 mL PBS, before 10 μL, 100 μL and the remaining volume were plated on media without 5-ala and appropriate antibiotics to select for transconjugants. Transposon insertion mutant colonies appeared up to five days after conjugations were performed. To obtain a pure culture of the transposon insertion mutants, colonies were restreaked at least three times.

### Screening for fluorescent colonies and fluorescence microscopy

For convenient and quick assessment of colony level fluorescence, a blue light gel reader was used for fluorescent proteins with emission wavelengths between 500-680 nm. Fluorescence microscopy was performed on a Zeiss AxioImager.M1 fluorescent widefield microscope equipped with Zeiss filter sets 38HE, 43HE, 46HE, and 47HE, (BP 470/40-FT 495-BP 525/50, BP 550/25-FT 570-BP 605/70, BP 500/25-FT 515-BP 535/30, and BP 436/25-FT 455-BP 480/40, respectively), an Axiocam 506, and the software Zeiss Zen 2.3. Single-cell fluorescence was analysed as described previously (Remus-Emsermann et al., 2016). In short, bacteria were mounted on an agarose slab (~1 mm thick, 1% agarose in milliq water) and samples were analysed using a Zeiss AxioImager.M1 at 1000 x magnification. Using ImageJ/Fiji (Schindelin et al., 2012).

### Arbitrary PCR

The transposon insertion of fluorescently (FP)-tagged bacterial strains was mapped using arbitrary PCR (Das et al., 2005). Briefly, a first PCR was performed using a mix of random primers Arb1, Arb2, and Arb3 containing an adapter oligo at the 5,-end and a primer targeting the transposon insertion (Table 2). Amplification was performed in a total volume of 25 μL containing: 15.05 μL ddH_2_O, 5 μL Phusion GC buffer, 1.25 μL 10 mM dNTP mix, 1.25 μL 10 μM arbitrary primers, 0.5 μL 10 μM ARB-RB-PCR1, 0.75 μL DMSO, 0.2 μL Phusion, and 1 μL DNA template. Cycling conditions were as follows: initial denaturation of 95 °C for 5 min; 6 cycles of 30 s at 95 °C for denaturation, 30 s at 30°C for annealing, and 1.5 min at 72 °C for extension; then, 30 cycles of 30 s at 95 °C for denaturation, 30 s at 45°C for annealing, 2 min at 72 °C for extension, and 5 min at 72 °C for a final extension. Then, PCR products were purified using the DNA concentrator kit (Zymo) following the manufacturer’s recommendations and eluted in 30 μL. A second PCR was performed using primers PCR2-AP and ARB-RB-PCR2. Amplification was performed in a total volume of 25 μL, containing: 12.8 μL ddH_2_O, 5 μL Phusion GC buffer, 1.25 μL 10 mM dNTP mix, 1 μL 10 μM arbitrary primers, 1 μL 10 μM ARB-RB-PCR1, 0.75 μL DMSO, 0.2 μL Phusion, and 3 μL of ten-fold dilution of purified PCR fragments. Cycling conditions were as follows: initial denaturation of 95 °C for 3 min; 30 cycles of 30 s at 95 °C for denaturation, 30 s at 52°C for annealing, 2 min at 72 °C for extension, and 5 min at 72 °C for a final extension. PCR products were then purified and concentrated using the Zymo Research DNA Clean & Concentrator^TM^-5 following the manufacturer’s recommendations and sequenced using Sanger’s sequencing (Macrogen). The sequencing results were mapped against the genome of the corresponding bacterial strain using Blast (Altschul et al., 1990) and a local database of draft genome sequences of the strains used in this study and the software Geneious prime (version 2020.1.2, Biomatters, Auckland).

## Results

We constructed a total of 23 plasmid vectors carrying different combinations of fluorescent proteins ranging from cyan to near-infrared fluorescence and antibiotic resistances including chloramphenicol, chloramphenicol and gentamicin, chloramphenicol and kanamycin, as well as chloramphenicol, kanamycin and erythromycin (Figure 1, Table 3). For fluorescent protein emissions and excitation spectra, please refer to Schlechter et al. (2018).

**Figure 1:**
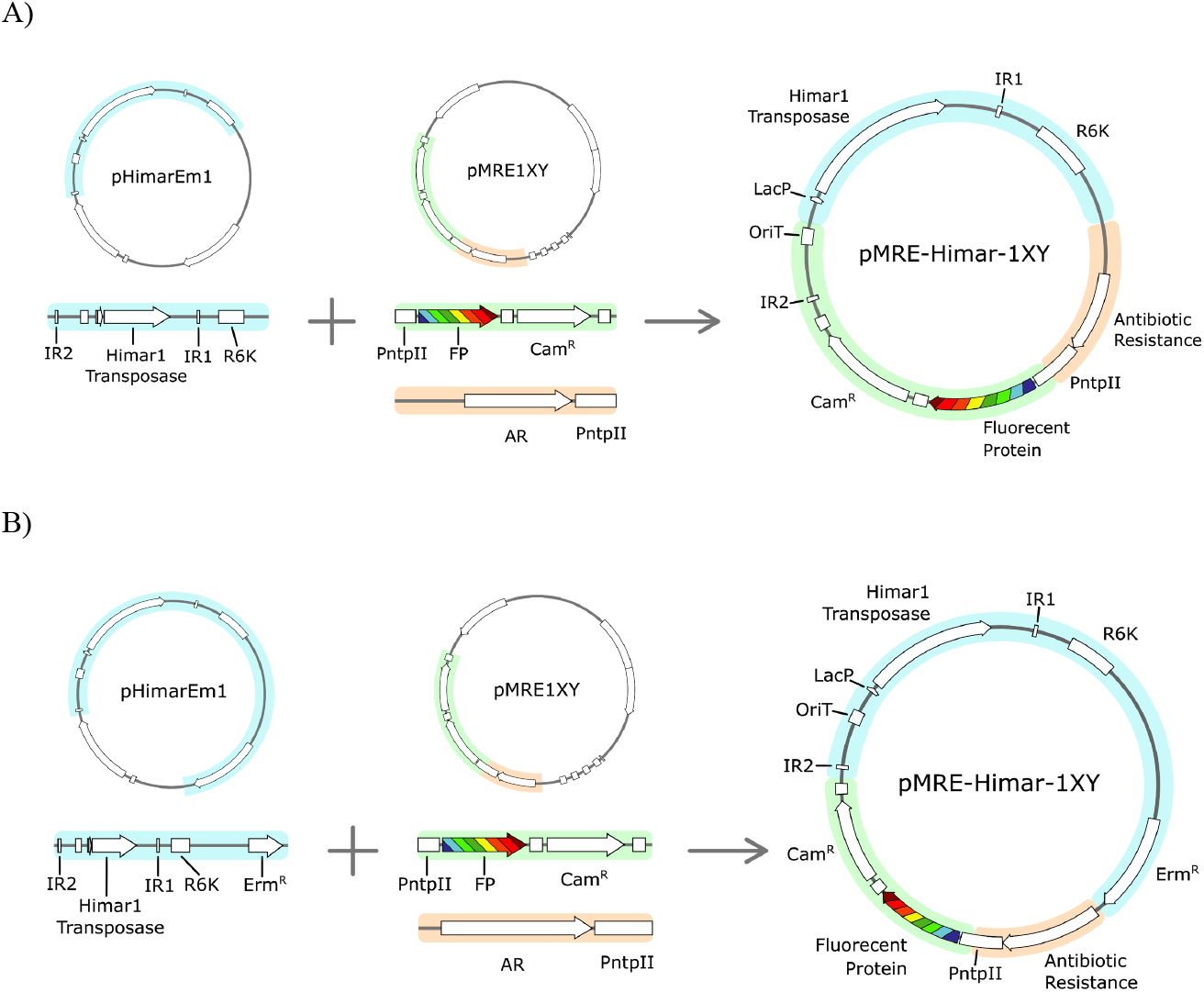
Overview of cloning procedures. A) to construct pMRE-Himar-13Y to pMRE-Himar-15Y, the pHimarEm1 backbone was amplified using PCR and Gibson assembly primers without the erythromycin resistance gene. Plasmids from the original Chromatic Bacteria series were PCR amplified in two separate reactions using Gibson assembly primers targeting the chloramphenicol resistance gene and the promoter driving the respective fluorescent protein gene expression and, where applicable, a secondary antibiotic resistance gene. These three fragments were joined using isothermal assembly. B) To produce pMRE-Himar-19Y plasmids, the procedure was similar as described in A), but in this case, the erythromycin resistance gene encoded on the pHimarEm1 plasmid was included when amplifying the backbone.

### Transposon delivery by conjugation

Using *E. coli* ST18 as a plasmid donor, conjugations were performed into the recipient strains in Table 1. It was possible to obtain transposon insertion mutants for four Proteobacterial strains as indicated in table 1, all mutants exhibited detectable fluorescent protein emission as determined using macroscopical observations at the single colonie level (data not shown) and at the single cell resolution as determined using widefield epifluorescence microscopy (Figure 2). The insertion sites for several independent insertion events were determined using arbitrary PCR and sequencing (Supplemental table 1). We were not able to obtain insertion mutants for the others strains listed in Table 1.

**Figure 2:**
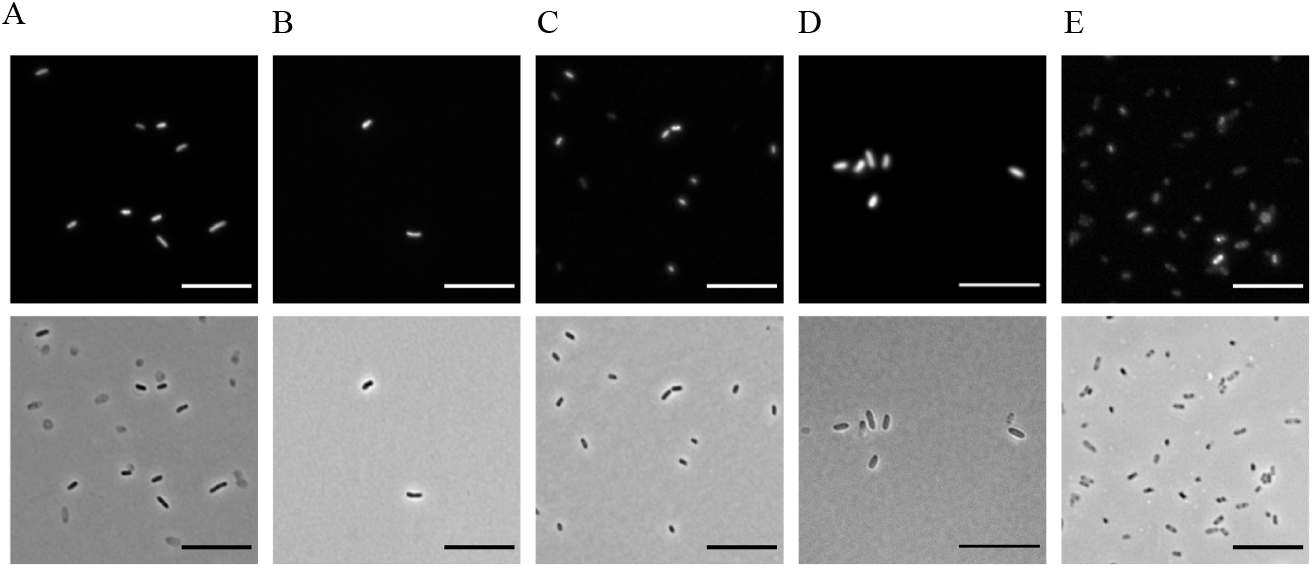
Fluorescence of recipients after stable integration of constitutively expressed fluorescent protein genes at the single cell resolution. Representative images of fluorescently-tagged bacterial strains. In all cases, fluorescence (upper panel) and phase contrast images (lower panel) are included. (A) *Acidovorax* sp. Leaf84::MRE-Himar-145. (B) *Sphingomonas* sp. Leaf357::MRE-Himar-145. (C) *Sphingomonas melonis* FR1::MRE-Himar-156; (D) *Pantoea eucalypti* 299R::MRE-Himar-145; (E) *Escherichia coli* ST18 (pMRE-Himar-156). Scale bar = 10 μm.

## Discussion

A range of new plasmids for the Chromatic Bacteria toolbox (Schlechter et al., 2018) based on R6K origin of replication plasmid backbone containing the Himar1 transposase was constructed. Additionally, a series of plasmids containing the Erm^R^ gene was constructed. A full list of plasmids constructed is included in table 3. The previous and novel antibiotic resistance gene combinations make the plasmids a versatile tool for bacterial genetic manipulation that accounts for different antibiotic resistances that might already be encoded by the recipient organism. Even though we constructed these additional new plasmids in *E. coli* S17-1 (Simon et al., 1983; Schlechter et al., 2018)‘ we have decided to clone them into *E. coli* ST18 (Thoma and Schobert, 2009). Similar to *E. coli* S17-1, *E. coli* ST18 allows conjugation into recipients, however, *E. coli* ST18 is lacking the *hemA* gene. The lack of *hemA* results in a strong auxotrophy and dependency of *E. coli* ST18 on exogenously provided 5-aminolevulinic acid. Thereby, there is no need to counterselect against the conjugation donor after conjugation by using minimal media or intrinsic antibiotic resistances of the host. The transposon mutants can be selected on their respective optimal complex medium with the addition of an antibiotic selecting for the transposon. Since the vector contains the R6K origin of replication, it only replicates if the λpir factor is present and expressed in the host, i.e. the plasmids are suicide vectors that are not able to replicate in their recipient.

Using the newly developed vectors, we were able to deliver Himar transposons randomly into bacterial recipients, similarly to the previously described pMRE-Tn5-transposon series (Schlechter et al., 2018). However, using the Himar1 transposase, we were able to genetically modify additional recipients that we previously failed to modify. In comparison to pMRE-Tn5-transposon series ‘ we were able to deliver himar transposons into *Acidovorax* sp. Leaf85, and *Sphingomonas* sp. Leaf357, while retaining the ability to deliver himar transposons into *S. melonis* Fr1 and *P. eucalypti* 299R. The enhanced spectrum of successful transposition can likely be accredited to the properties of the Himar1 transposase that does not require additional host cofactors to function (Lampe et al., 1996). However, we were not successful in conjugating the plasmids and insert the transposons into the genomes of non-model Gram-positive bacterial strains including *Aeromicrobium* sp. Leaf245, *Agreia* sp. Leaf 335, *Arthrobacter* sp. Leaf145, *Microbacterium* sp. Leaf320, *Microbacterium* sp. Leaf 347, *Plantibacter* sp. Leaf1, *Rathayibacter* sp. Leaf296, *Rhodococcus* sp. Leaf225, or *Williamsia* sp. Leaf354.

The R6K origin of replication is present in the transposon, which allows, next to the above described arbitrary PCR, to determine the insertion site of the transposon into the recipient’s genome. To that end, the genome of the recipient can be isolated, digested with a rare cutting enzyme that does not cut in the transposon sequence, such as KpnI, and then re-ligated and transformed into *E. coli* S17-1 or another λpir factor expressing *E. coli* cloning host (Metcalf et al., 1994). Due to the R6K origin of replication contained and the antibiotic resistances located in the transposon, this will result in functional plasmids. These plasmids can then be sequenced using a sequencing primer (Table 2) to determine the sequence of the insertions site. This process is significantly more time intensive and less cost effective compared to the arbitrary PCR described above, but might be advantageous in cases where the arbitrary PCR does not yield results of sufficient quality since the procedure yields results with a better signal to noise ratio compared to arbitrary PCRs.

As described in Schlechter et al. 2018, the here constructed plasmids allow for convenient fluorescent tagging of many environmental bacteria and cover a different host range compared to the previously described vectors. Fluorescent protein tags are the prerequisite for many experimental studies that follow different populations simultaneously, identify focal populations in complex environments, or to follow the behaviour of individual cells (Wang et al., 2010; Diard et al., 2013; Ledermann et al., 2015; Remus-Emsermann and Schlechter, 2018). Currently, many delivery systems, including the first versions of the chromatic bacteria, function almost exclusively in Proteobacteria, the Himar transposons have been shown to have a wider range of activity. Thereby the here introduced plasmids can serve as a one fits many solution for tagging proteobacteria.

## Conclusion

The here-described plasmids extend the previously constructed Chromatic bacteria toolbox host range and utility. All plasmids and plasmid sequences will be made available through the not for profit service addgene.org upon peer reviewed publication (Addgene plasmids numbers to be determined).

## Acknowledgement

The authors thank Mark Silby and Lucy McCully (University of Massachusetts, Dartmouth) for the kind gift of pHimarEm1. This project was supported by the New Zealand Tertiary Education Commission CoRE grant to the Bio-Protection Research Centre and a Marsden Fast-Start grant to MR-E (UOC1704) and additional funding by Callaghan Innovation and the Bioprotection Research Core (BPRC) is acknowledged. CS is supported by a UC Masters scholarship and a Bioprotection Research Core summer scholarship. RS is supported by a New Zealand International Doctoral Research Scholarship (NZIDRS) and a University of Canterbury Doctoral Scholarship.

