## Supplementary table 1 for "Chromatic bacteria v.2 - A Himar1 transposon based delivery vector to extend the host range of a toolbox to fluorescently tag bacteria"

**Supplementary material**

**Supplemental table 1**. Transposon insertion sites

| Strain name | Transposon insertion flanking region sequence | Region hit |
|---|---|---|
| *Sphingomonas* sp. Leaf357::MRE-Himar-145/1 | AGGGGCTCGCAGTCGATTTACCGGTTCGCATGATCGTAACCGCACAGGGGAAGGAAACATGGGCTCTCTTCCGCCAGCGCGGTGGGATGTACCCTGAG | Beta-hexosaminidase CDS |
| *Sphingomonas* sp. Leaf357::MRE-Himar-145/2 | TGTTATAACCCGGGGCCCAGAAGCGCGCGAGGTAGTCTTTGAATGGATACATGGGCAGATATGCGATAACGCCGTCGAGCTTCCGGTTGGCGACGTCTCAGTCCGCGTCCATGACGACCCCGAGCGTGT | Hypothetical protein |
| *Sphingomonas* sp. Leaf357::MRE-Himar-145/4 | CTCGGCGCGCAGGCCAATCTGTGGGCCGAATATATCGTGACGCCCACCGAATCCCAACATGCGCTGTTCCCGCGCGTCGACGCGCTGGCCGAGATCGCCTG | Hypothetical protein |
| *Acidovorax* sp. Leaf84::MRE-Himar-145/2 | AGTCAACATCGAAAAGCTCGGAGACTATGTGAATCGCTAT-GGCGTCAATAGCTTTTTCGACGCATCCGATGATGCCCATC | Intergenic region |
| *Acidovorax* sp. Leaf84::MRE-Himar-145/5 | ATCTGATCTTCAGACAGTCTGTCGGTAGCTCCCTCGCGCCTTGCAGAGCAGATGATGTGTTCCCCTTGAAAACGCCCTTGACATCATGCACCTCGACG | Hypothetical protein |
| *Acidovorax* sp. Leaf84::MRE-Himar-145/6 | TAATCGGTGGATGGTAAATAGATAGGAAATTTATCACTGTGTTTCATAACAGGTTG | Intergenic region |
